# Protection against necrotizing enterocolitis by fecal filtrate transfer requires an active donor virome

**DOI:** 10.1101/2024.09.23.614450

**Authors:** Malene Roed Spiegelhauer, Simone Margaard Offersen, Xiaotian Mao, Michela Gambino, Dennis Sandris Nielsen, Duc Ninh Nguyen, Anders Brunse

## Abstract

Necrotizing enterocolitis (NEC) remains a frequent catastrophic disease in preterm infants, but fecal filtrate transfer (FFT) has been identified as a promising prophylactic therapy in preclinical studies. This study examined the importance of the FFT virome viability on gut colonization and NEC occurrence. We established an ultraviolet irradiation-based viral inactivation protocol and demonstrated total loss of infectivity of a viral mock community. Using this protocol, we inactivated an aliquot of sterile-filtered donor feces and compared the response in preterm piglets subjected to experimental NEC induction. Gut pathology and barrier properties were assessed, and bacterial and viral compositions were determined by 16S rRNA amplicon and viral metagenomics sequencing, respectively. Native FFT decreased NEC severity and proinflammatory cytokines, but inactivated FFT (iFFT) completely abolished these effects. Mild side effects in the form of diarrhea manifested earlier in recipients of native FFT than iFFT or controls. A distinct gut colonization pattern of increased viral heterogeneity increased bacterial homogeneity and reduction in pathobionts like *Clostridium perfringens* and *Escherichia* was observed in the group receiving native FFT, but not in the iFFT group. The present study uncovered a clear distinction between active and inactivated transferred viromes in the ability to modulate gut colonization after preterm birth and decrease NEC. FFT efficacy is potentially driven by active bacteriophages targeting pathogenic bacteria.

## Introduction

Necrotizing enterocolitis (NEC) is a serious gastrointestinal disease with an incidence of 2-10% of infants born before week 32 of gestation), and up to 17% of infants born with very low birth weight (<1500 grams) [1], [2]. The mortality of infants with NEC is inversely related to birth weight and increases from 16% in infants below 1500 grams to 42% in the smallest infants weighing less than 750 grams [3]. NEC is characterized by excessive intestinal inflammation and tissue necrosis [4], and life-saving surgical resection of the diseased intestine is often necessary as tissue injury has already manifested at the onset of clinical symptoms [5].

Gut microbiome dysbiosis, including loss of bacterial diversity and overgrowth of single species, constitutes a risk for NEC development [6], [7]. As the maturational trajectory of the preterm gut microbiome is prolonged compared to full-term infants [8], [9], they consequently harbor otherwise transient bacteria with pathogenic potential for longer periods. The interval in which these bacteria dominate the preterm gut seems to coincide with the time at which the risk of NEC is highest [10].

No single bacterial species has been consistently associated with NEC, but several studies have described an alteration of the gut microbiome prior to NEC onset, characterized by an increased relative abundance of *Pseudomonadota* members such as *Klebsiella* spp., and decreased relative abundance of *Bacillota* and an overall reduced bacterial diversity [4], [8], [11], [12], [13]. In addition to bacteria, the gut microbiome contains a complex population of bacteriophages (phages), viruses that exclusively infect bacteria in a host-specific manner. Phages form a central part of the human intestinal virome, where they occupy the gut from the first week of life [14], [15], also in preterm infants [16], [17]. A recent study illustrated the relationship between the virome and NEC development, by shifts in the preterm gut virome before the onset of NEC, and an increase of specific phage signatures related to bacteria like *Escherichia* or *Streptococcus* [18].

We have previously tested fecal microbiota transplantation (FMT) in a preterm piglet model of NEC [19], [20]. The FMT effectively reduced NEC but also increased the risk of sepsis from translocating bacteria, and combination with antibiotics eliminated the protective effect of FMT. Instead, administration of a sterile fecal filtrate (FFT), where bacteria were removed by filtration (0.45 µm filter), fully protected against NEC without causing sepsis or adverse effects [21]. FFT also reduced the abundance of *Enterobacteriaceae* in close proximity to the ileal mucosa and increased gut virome diversity. The detailed mechanism behind this effect is unclear, but we suggest that it is likely due to the bacteriolytic effect of transferred phages.

This study aimed to assess if the efficacy of FFT for protection against NEC depends on the phages being infective. We addressed this by inactivating the virome in the donor fecal filtrate through irradiation at the UV-C spectrum (200-280 nm), which is known to cause viral inactivation through damage to nucleic acid polymers [22]. We then transferred native or UV-inactivated donor fecal filtrate or control vehicle into NEC-susceptible preterm piglets and recorded gastrointestinal pathological responses and microbiome composition.

## Materials and Methods

### Fecal filtrate preparation

The donor fecal filtrate was prepared as previously described [21]. Briefly, colon content was collected from four 10-day-old healthy piglets from four different herds after euthanasia. The contents were pooled, diluted 1:1 in sterile 20% glycerol, and stored at 4°C. The following day, the pool was further diluted 1:10 in SM buffer (200 mM NaCl, 10 mM MgSO_4_, 50 mM Tris-HCl, pH 7.5) and centrifuged at 5000 x *g* for 30 minutes at 4 °C. The supernatant was passed through a syringe filter with a pore size of 0.45 µm (Minisart Syringe Filter Polyethersulfone (PES), Sartorius).

### Ultraviolet irradiation

Viral susceptibility to nucleic acid damage was assessed by subjecting six phages representing different phage types and host ranges [23], to increasing ultraviolet (UV) irradiation exposure time (265 nm wavelength, UV LED Collimated Beam Device PearlLab Beam^TM^, Aquisense Technologies). This was based on a protocol developed by Vitzilaiou et al. [24]. The six phages were subjected to UV-irradiation for 0, 10, 20 or 30 minutes in titers between 10^3^-10^6^ plaque-forming units (PFU)/ml to identify a minimum dose that would eliminate the infectivity of all phages (Suppl. Table S1). The infective titer of UV-irradiated phages was estimated by plaque assay as previously described [25]. Next, a portion of the donor fecal filtrate was UV-irradiated for 45 minutes, receiving a total dose of 232 mJ/cm^2^ to ensure total virus inactivation. The native fecal filtrate and the UV-inactivated fecal filtrate were divided into aliquots and stored at -80 °C while ensuring they were not exposed to sunlight before use.

### Virus particle enumeration

The concentrations of virus-like particles (VLPs) in the fecal filtrates were estimated following a previously described protocol [26]. In short, the filtrates were stained with SYBR Gold (Thermo Fisher Scientific), blotted on 0.02 µm filter membranes (Anodisc, Whatman) and mounted on glass slides. VLPs were visualized with an epifluorescence microscope and a Photometrics CoolSNAP camera at 1000x magnification using a 490 nm filter block. Ten representative pictures were taken of each filtrate and the number of VLPs was estimated by ImageJ software [27] with a lower detection limit of 2.5 × 10^5^ VLPs/ml filtrate.

### Animal housing and clinical evaluation

All experimental procedures were approved by the Danish Animal Experiments Inspectorate (license no. 2020-15-0201-00520) and followed ethical guidelines of laboratory animal care. Thirty-nine piglets were delivered by caesarean section at 90% gestation from three healthy, conventionally bred sows (Landrace x Yorkshire x Duroc). The experiment was carried out alongside another study [28], with shared control groups (CON and FFT groups) to reduce the total number of experimental animals used. All animals received passive immunization by infusion with sow’s plasma (16 ml/kg) on the first day of life. Enteral nutrition was administered to all animals through an oral catheter every 3 hours (the daily doses were 24, 40, 64, 96 and 96 ml/kg^0.75^/day from day one to five, with the same formula composition as in [28]), together with continuous infusion of parenteral nutrition (96, 96, 72, 48, 48 ml/kg^0.75^/day from day one to five, Kabiven 3-chamber bags, Fresenius-Kabi) as described previously [19].

### Administration of treatments

Animals were stratified by sex and birthweight and randomly allocated to three treatment groups receiving either native donor fecal filtrate (FFT), UV-inactivated donor fecal filtrate (iFFT) or SM buffer as control (CON). The animals each received 1.0 ml of the respective treatments twice per day for the first two study days.

### Clinical evaluation and euthanasia

Animals were regularly monitored by experienced staff and clinically assessed every third hour at a minimum. The animals were weighed daily, and clinical status and stool consistency were reported every 12 hours. Clinically affected animals were monitored closely and administered pain relief if possible. Animals showing clinical signs of NEC (defined as abdominal distension, reduced activity, cyanosis, and discoloration), or signs of discomfort that could not be relieved, were euthanized and necropsied to identify the cause of illness. On study day five, animals were deeply anesthetized and euthanized by intracardiac injection of Euthanimal (Sodium pentobarbital, 400 mg/ml, Alfasan).

### Gastrointestinal pathology

Pathological lesions in the stomach, small intestine and colon were assessed in a blinded manner by a cumulative scoring system based on the distribution of hyperemia, hemorrhage, pneumatosis intestinalis, and necrosis [29]. The diagnosis of NEC was defined by moderate hemorrhage or pneumatosis, or the presence of necrosis.

### Sampling procedure

Abdominal organs were sampled and weighed. A distal part of the small intestine and the colon were collected in Carnoy’s solution (Ampliqon). Duplicate parts of the small intestine and colon with lesions were collected in cryotubes for freezing and tissue cassettes for immersion in 4% paraformaldehyde. One colon lesion with content was collected in a cryotube. A 10 cm section of distal ileum was excised, rinsed with sterile saline and cut open longitudinally. The mucosa was then scraped off and collected in a cryotube. All collected cryotubes were immediately submerged in liquid nitrogen and subsequently stored at -80 °C. Bone marrow was aseptically sampled from the left femoral bone and stored at 4 °C.

### Histopathology and mucin proportion

Formaldehyde-fixed small intestine and colon were embedded in paraffin and cut into several 3-5 µm sections for different microscopy analyses. *<u>H&E</u>.* Sections were stained with hematoxylin and eosin (H&E), and the severity of histopathologic changes was evaluated microscopically using a scoring system defined previously [29].

#### AB-PAS

Sections were stained with Alcian Blue and periodic acid-Schiff (AB-PAS) to visualize goblet cells. Four digital images per tissue section were captured at 10x magnification using a DM2500 light microscope system with a camera attached (Leica Microsystems, Wetzlar, Germany), and the relative area of mucin-containing goblet cells was estimated using Fiji image analysis software [30].

### Fluorescent in situ hybridization (FISH)

Tissue collected in Carnoy’s solution were transferred to ethanol, embedded in paraffin, and cut into 5 µm sections. The sections were stained with fluorescent DNA hybridization probes specific for all bacteria (EUB338, 5’ GCTGCCTCCCGTAGGAGT ’3, Alexa-Fluor-555 labelled, Eurofins Genomics) and *Pseudomonadota* specifically (GAM42A, 5’ GCCTTCCCACATCGTTT ‘3, Alexa-Fluor-647 labelled, Eurofins Genomics) following a previously described protocol [31]. Autofluorescence was removed with TrueView Autofluorescence Quenching Kit with DAPI (Vector Laboratories) and cover slides were mounted with 40 µl VectaShield Vibrance Antifade Mounting Medium with DAPI (Vector Laboratories). The stained sections were evaluated by confocal fluorescence microscopy (ZEISS LSM 900, software Zeiss Zen Blue 3) at 20x magnification in a blinded manner and assigned a semi-quantitative bacterial density score: 0 (no bacteria), 1 (low density in single areas), 2 (low density across tissue), or 3 (high density across tissue).

### Gut cytokines and enzymes

Frozen small intestine and colon tissue were dissociated in 1% Triton-X solution (GentleMACS dissociator, Miltenyi Biotec). The levels of the inflammatory cytokines interleukin (IL) 1β and IL-8 were measured by enzyme-linked immunosorbent assays (ELISA) (Porcine-specific DuoSet kits, R&D Systems). The activity of brush border enzymes Aminopeptidase A (ApA), Aminopeptidase N (ApN), Dipeptidyl peptidase IV (DPPIV), lactase, maltase, and sucrose were measured from dissociated ileal mucosa as described previously [32].

### Bone marrow bacteriology

Collected bone marrow was homogenized in 2 ml sterile 0.9% saline (Merck), diluted, and triplicates of 20 µl were cultured aerobically at 37 °C (LB Agar Lennox, BD Beckton Dickinson and Company). Bacteria were enumerated as the number of colony-forming units (CFU) per gram tissue with a lower detection limit of 16.6 CFU/ml homogenate. The bacterial isolates were identified with MALDI-TOF MS (Bruker).

### Cell culture and challenge

THP-1 monocytes were cultured in RPMI 1640 Glutamax (Gibco, ThermoFisher Scientific) supplemented with 10% heat-inactivated fetal bovine serum (FBS, Gibco, ThermoFisher Scientific) and 25 µg/ml Gentamicin (Gibco, ThermoFisher Scientific) in humidified cell incubators at 37 °C with 5% carbon dioxide. The cells were cultured in concentrations between 5 × 10^5^ to 1 × 10^6^ cells/ml, seeded and differentiated into macrophages following a modified protocol [33], [34]. Briefly, a solution of 1 × 10^6^ cells/ml supplemented with 5 ng/ml phorbol 12-myristate 13-acetate (PMA, ThermoFisher Scientific) was seeded in each well of a 12-well cell culture plate (TPP) and incubated for 48 hours. The medium was replaced with RPMI 1640 Glutamax supplemented with 10% FBS (without PMA) one hour before adding either 100 µl FFT, iFFT or SM buffer (as control of unchallenged cells). The plate was centrifuged for 5 minutes at 300 rpm and incubated for 20 hours. The challenges were performed in triplicates using cell passages 17, 19 and 20.

### Relative quantification of gene expression

Total RNA was extracted from the THP-1 derived macrophages (Qiagen RNeasy Mini Kit, Qiagen) and reverse transcribed to cDNA (High-Capacity cDNA Reverse Transcription Kit, Applied Biosystems and MultiGene Optimax Thermocycler, LabNet International). Quantitative PCR (qPCR) was performed in duplicate reactions of 10 µl volumes (LightCycler 480SYBR Green Master, Roche Holding AG) using 1× SYBR green, 0.4 µM forward primer and 0.4 µM reverse primer (TAG Copenhagen, Denmark). Primer sequences are listed in Suppl. Table S2. The qPCR cycling conditions were as follows: initial denaturation at 95 °C for 15 minutes, 45 cycles of amplification (94 °C for 15 seconds, 58 °C for 30 seconds, 72 °C for 30 seconds), and final elongation at 72 °C for 10 minutes, with measure of fluorescence after each cycle (LightCycler 480 II and LightCycler 480 Software release 1.5.1.62 SP3, Roche). The relative gene expression was calculated from the geometric average of the reference genes *PGK1* and *ACTB*.

### Microbiome analysis

Native FFT and iFFT inoculums were thawed, suspended in 5 ml SM buffer and homogenized at 300 rpm for 10 minutes. Sampled colon lesions with content were homogenized in 5 ml SM buffer (Stomacher 80 Biomaster Lab Blender, Seward) for two minutes and centrifuged at 5000 x *g* at 4 °C for 30 minutes. The pellets were used for bacterial 16S rRNA gene amplicon sequencing, while the supernatants were saved for metavirome sequencing.

#### Bacterial DNA extraction, 16S rRNA gene amplicon sequencing and data processing

140 µl of the fecal filtrates (FFT and iFFT) were pretreated with 5 units of Pierce Universal Nuclease (ThermoFisher Scientific) for 15 min at room temperature before the DNA extraction. Bacterial DNA was extracted from the colon homogenate pellets using the DNeasy PowerSoil Pro Kit (Qiagen) following the manufacturer’s protocol. The concentration of extracted DNA was measured on Qubit 1x dsDNA High Sensitivity kit on Varioskan Flash (Thermo Fisher Scientific) and stored at -80 °C. A near-full length of 16S ribosomal RNA (rRNA) gene amplicon was sequenced with GridION (Oxford Nanopore Technologies) [35] and the data were collected with the software MinKNOW (v22.10.7, Oxford Nanopore Technologies). Raw FAST5 data were base-called to FASTQ format (Oxford Nanopore Technologies) with the Guppy v6.2.8 base-calling toolkit. Abundance tables were generated from the raw FASTQ-files with the Long Amplicon Consensus Analysis pipeline [36]. The quality-corrected reads were assigned taxonomy using the SILVA database.

#### Viral DNA/RNA extraction, metavirome sequencing and data processing

The fecal inoculum and recipient fecal supernatants were passed through a 0.45 µm PRE membrane filter (Filtropur, Sarstedt) and concentrated by centrifugation at 1500 x *g* at 4 °C using ultrafiltration devices with a filter cut-off at 100 kDa (Centrisart, Sartorius) [37]. Before viral DNA/RNA extraction, 140 µl of the collected supernatant, the FFT and iFFT inoculum and the SM inoculum were treated with 5 units of Pierce Universal Nuclease (ThermoFisher Scientific) for 15 minutes at room temperature. Viral DNA/RNA was extracted with the Viral RNA Mini Kit (Qiagen) as described previously [38]. Reverse transcription was performed using Superscript VILO Master Mix (Invitrogen, Thermo Fisher) according to the manufacturer’s instructions, followed by purification with DNeasy Blood and Tissue Kit (Qiagen) only following steps 3-8 of the protocol. To include ssDNA viruses, multiple displacement amplification was used using the GenomiPhi V3 DNA Amplification Kit (Cytiva). The sequencing library was prepared with the Nextera XT Kit as previously described [38] and sequenced with the NoveSeq platform (NovoGene). Viral operational taxonomic units (OTU) tables were prepared using a modified version of the Vapline v2.0 [39]. In brief, the raw reads were quality-controlled, trimmed and dereplicated with Trimmomatic, BBmap and Seqkit [40], [41], [42]. Reads were paired and assembled within each sample with SPAdes v3.13.0 using both paired and unpaired reads [43]. Contigs shorter than 2200 base pairs were removed. The remaining contigs were analyzed with CheckV, VirSorter2 and VIBRANT [44], [45], [46]. Contigs that were identified as viral in at least one of the three programs were classified as viral contigs and included in the analysis. The viral contigs were assigned to a taxonomy by aligning to a custom database with mmseqs2 based on the VOGDB (v217), and host prediction was performed with iPHoP [47]. Viral OTU tables were prepared by assigning raw reads from each sample to viral contigs with SAMtools [48].

### Statistical analyses

All statistical analyses were performed using RStudio version 2023.03.1. Figures showing clinical or cell results were created using GraphPad Prism version 9.5.1 or RStudio version 2023.03.1. Figures showing the microbiome results were prepared with the R-package “ggplot2”, unless directly specified [49].

Continuous data were compared using linear models with group and sex as fixed covariates and litter as random covariate, followed by Tukey pairwise comparison of groups. Mean time for diarrhea onset was evaluated by Cox regression analysis. The normalized bodyweights were compared using a linear mixed model with unstructured covariance pattern and group, sex, litter and birth weight as covariates. NEC scores were analyzed using ordinal logistic regression with group, sex, birthweight and litter as covariates, followed by Chi-squared test for likelihood ratio, least squared means and Benjamini Hochberg p-value adjustment. Categorical and non-parametric continuous data were analyzed with Kruskal-Wallis test followed by Dunn pairwise comparison of groups and Benjamini Hochberg p-value adjustment. Binary data were compared with Fisher’s exact test. Cell gene expression was log2-transformed and compared in a linear model with group and replicate as covariates, followed by Tukey pairwise comparison of groups. For bacterial and viral analyses, decontamination was performed with the R-package “decontam” using a threshold of 0.5 [50]. The R-package MetagenomeSeq was used to perform cumulative sum scaling (CSS) normalization [51]. The Adonis function in the R-package “vegan” contained the PERMANOVA adjusted by False Discovery Rate (FDR) to compare beta diversity [52]. Alpha diversity and Bray-Curtis dissimilarity were compared within groups with the Wilcoxon test adjusted by FDR. Differentially enriched microorganisms at summarized species level (bacteria) and OTU (viruses) were compared with DESeq2 [53], prepared in volcano plots using the R-package “EnhancedVolcano” with a p-value cut-off at 0.05 and log2 (fold change) cut-off at 0.6 [54]. The significance level was set to a probability value of less than 0.05.

## Results

### Ultraviolet irradiation is suited for universal phage inactivation

The donor fecal filtrate (FFT) was prepared using pooled colon content from four 10-day-old healthy-appearing farm piglets, according to a previously published protocol [21] (Fig. 1A). We initially assessed the UV-exposure time sufficient for viral inactivation using a mock community of six phages [23]. The phage infectivity decreased with UV exposure, and 30 minutes of irradiation was sufficient to inactivate all six phages (Fig. 1B, Suppl. Table S1). We then prepared an inactivated fecal filtrate (iFFT) by subjecting an aliquot of the fecal filtrate to UV-irradiation for 45 minutes (Fig. 1A).

**Figure 1.**
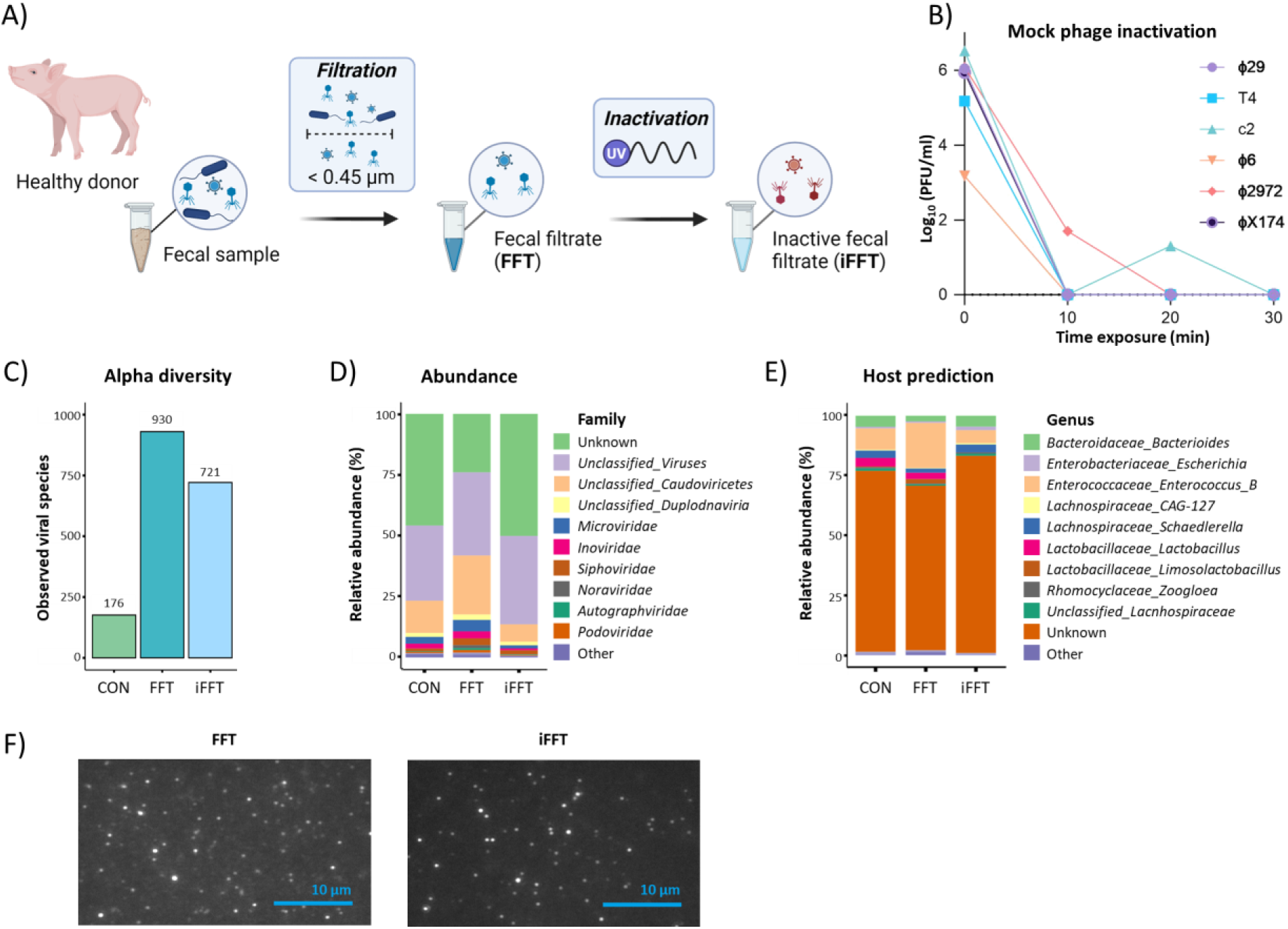
Donor fecal filtrate preparation, UV-irradiation and metagenomics sequencing. A) Preparation of donor fecal filtrate treatments. B) Mock phage community titers following UV-irradiation. C) Number of observed viral species. D) Virome composition. E) Predicted bacterial hosts. F) Visualization of virus-like particles (VLP) at 1000x magnification by fluorescence staining microscopy. PFU: Plaque-Forming Units. Panel A was created with BioRender.com.

The FFT, iFFT and sterile buffer (CON) were subjected to metavirome sequencing. Despite originating from the same batch, the native FFT and iFFT differed marginally in terms of numbers of observed viral species, although both were substantially higher than the background diversity of sterile buffer (Fig. 1C). Viral taxonomic composition and bacterial host prediction also differed between the filtrates (Fig. 1D-E). We suspect that the damage to viral nucleic acids caused by UV-irradiation might have affected the DNA amplification and viral assembly step in the metagenomics analysis pipeline. Nevertheless, the densities of virus-like particles estimated by fluorescence microscopy were similar between the filtrates (FFT: 3.38 x 10^9^ VLP/ml, iFFT: 2.36×10^9^ VLP/ml), which suggests that the virions remained intact (Fig. 1F).

### Reduction of intestinal pathology by fecal filtrate transfer requires a viable virome

Subsequently, we compared the *in vivo* effects of native fecal filtrate (FFT) and inactivated fecal filtrate (iFFT) using cesarean-delivered preterm piglets. In this animal model, formula feeding provokes spontaneous NEC-like pathology comparable to the pathology observed in preterm infants [55]. The study included thirty-nine piglets from three litters randomly allocated to receive FFT, iFFT or sterile buffer (CON) administered orally twice daily over the first two study days (Fig. 2A). The animals were sacrificed on study day five, where the gastrointestinal tract was subjected to pathological evaluation using a validated scoring system [29] (Fig. 2B). Seven animals were euthanized before the end of the study but were included in the data analysis: one CON, two FFT and two iFFT animals with suspected NEC, one FFT animal with respiratory distress and one iFFT animal with other complications. Additionally, one iFFT animal with very early onset NEC was euthanized before end of study and excluded from further analysis due to lack of detailed necropsy record.

**Figure 2.**
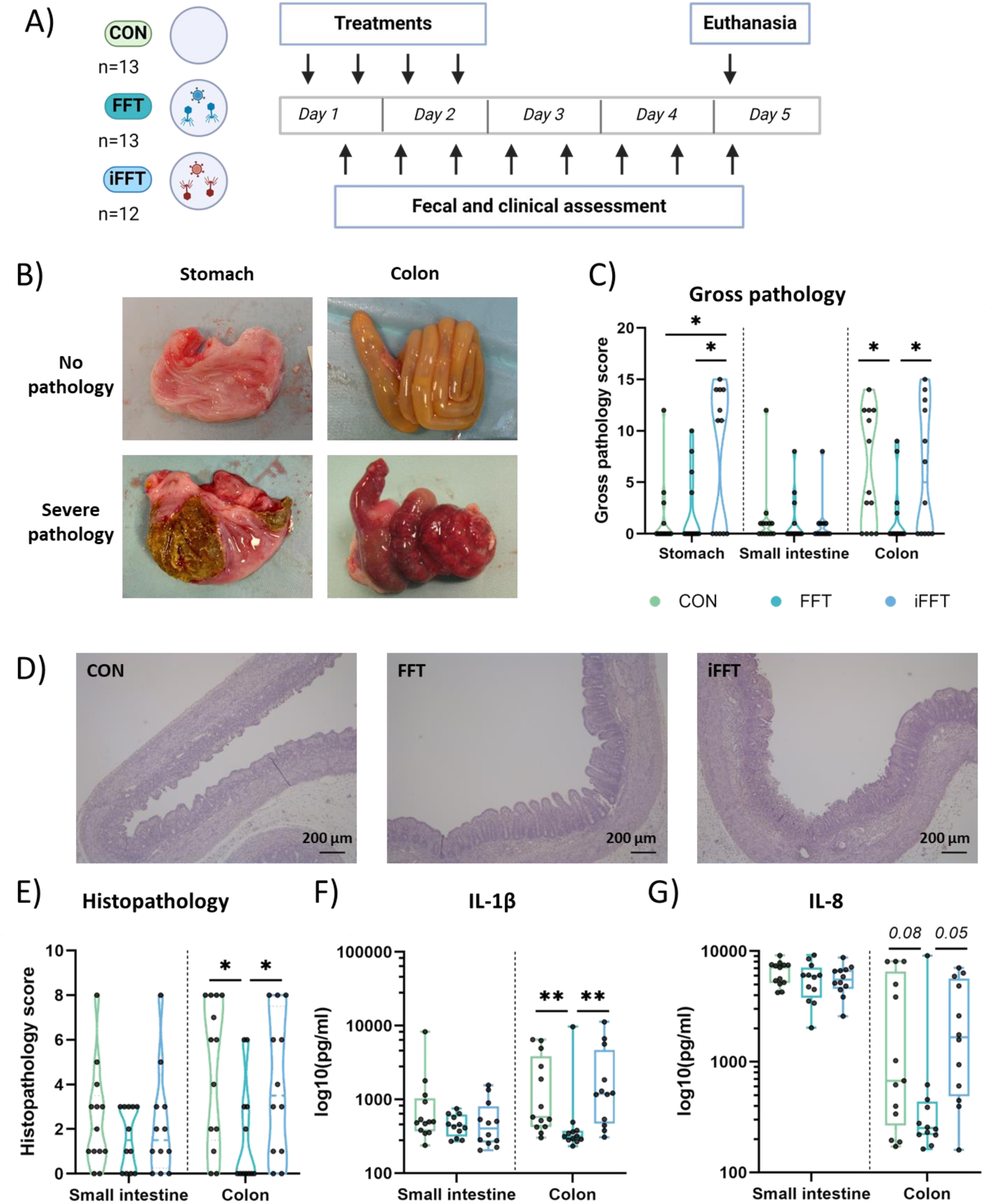
Reduced pathology score and intestinal inflammation was only observed in animals receiving an active fecal filtrate. A) Animal study design. B) Example of stomach and colon with no pathology (score 0) or severe pathology (score 12 and 15, respectively). C) Gross pathology scores. D) Representative hematoxylin & eosin (H&E) stain images of colon tissue, scale: 200µm. E) Histopathology scores. F) Interleukin (IL) 1β concentration (pg/ml). G) IL**-**8 tissue concentration (pg/ml). The continuous data is presented as boxplots with median and interquartile range, ordinal data is presented as violin plots. Significance levels are * p<0.05, ** p<0.01. n= 12-13 animals per group. Panel A was created with BioRender.com.

Small intestinal lesions occurred infrequently and similarly among the groups, whereas significantly reduced colonic gross pathology scores were observed in FFT recipients relative to both controls and iFFT recipients (Fig. 2C). Only the iFFT recipient group showed severe stomach pathology (Fig. 2C).

For validation purposes we stained paraffin-embedded sections of the small intestine and colon with hematoxylin & eosin and graded the severity of histopathological changes using an established method [29]. Again, we found that small intestinal changes were similar among groups, while the colonic histopathological severity was reduced only in the FFT recipient group, corroborating the gross pathological observations (Fig. 2D-E). We proceeded by measuring the concentrations of IL-1β and IL-8 in lesion areas of small intestine and colon tissue. Animals receiving FFT showed reduced colonic levels of interleukins IL-1β and IL-8 compared to the CON (p=0.007 and p=0.08) and iFFT (p=0.004 and p=0.05) groups, indicating a lower degree of tissue inflammation, whereas small intestinal cytokine levels remained unchanged among the groups (Fig. 2F-G). Taken together, these data clearly demonstrated the beneficial effects of FFT on intestinal pathology and inflammation which were eliminated by prior inactivation of the FFT. This supports our notion that the therapeutic effects of FFT are dependent on virome viability.

### Fecal filtrate transfer provides minimal changes to intestinal barrier, but uniform translocation of bacteria

To further investigate events occurring at the gut microbe-host interface, we evaluated goblet cell mucin density, bacterial adherence and translocation. Exhaustion of goblet cell mucin stores compromising barrier function is among the earliest changes to epithelial integrity in NEC pathogenesis [29], most likely in response to bacterial stimulation and limited *de novo* mucin synthesis. Initially, we stained small intestinal and colonic tissue sections with Alcian blue & periodic acid-Schiff (AB-PAS) to visualize acidic and neutral mucins and quantified the relative mucin area by image analysis. Small intestinal and colonic goblet cell mucin density appeared similar among the groups (Fig. 3A,C left panel).

**Figure 3.**
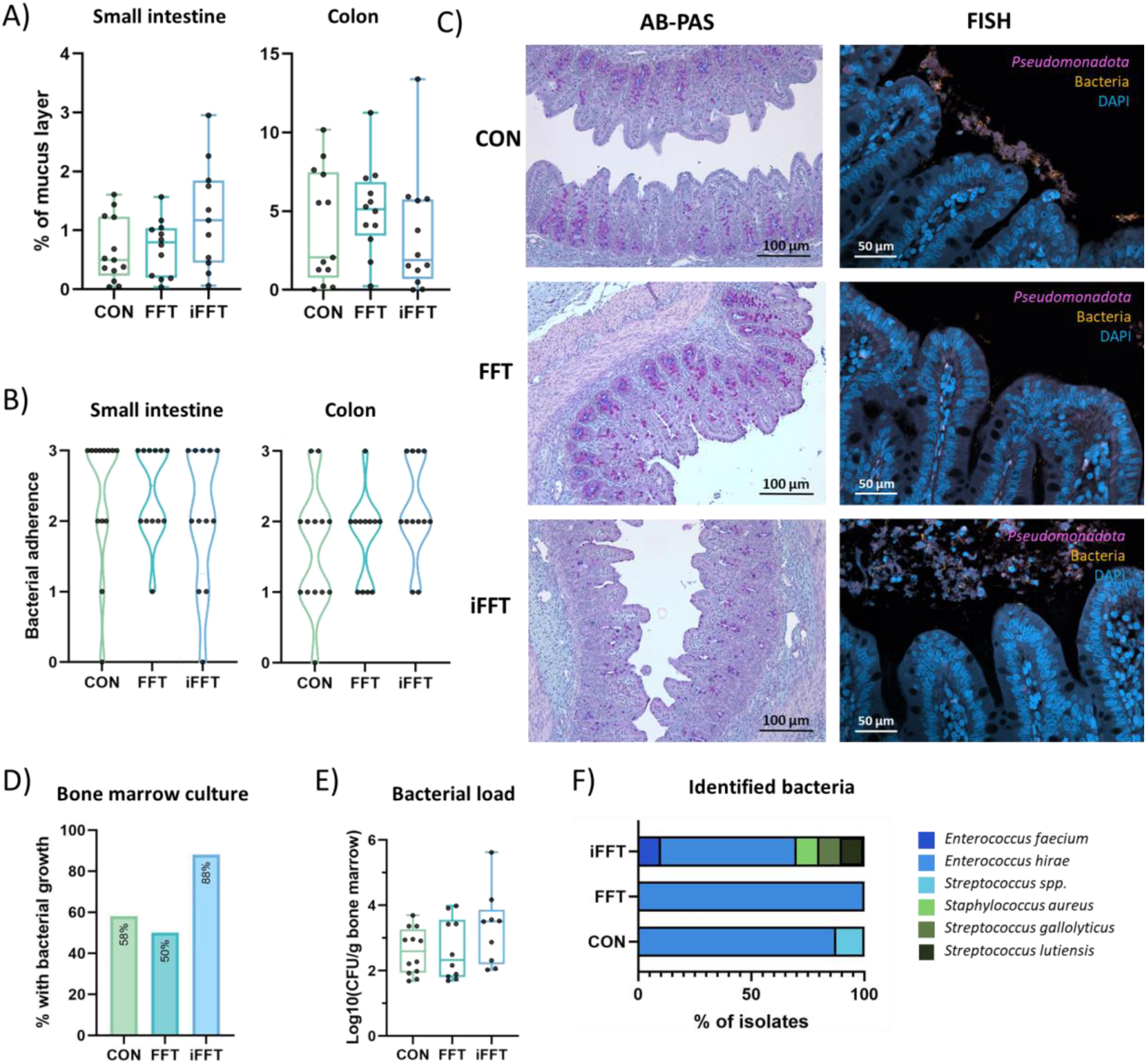
Minimal changes in gut barrier properties provided by either the native or inactivated fecal filtrates. A) Small intestine and colon goblet cell density. B) Score of bacterial adherence in small intestine and colon, ranging from 1-4. C) Left: Representative pictures of colon AB-PAS stain for goblet cell count. Right: Representative pictures of fluorescence microscopy for visualization of adherent bacteria. D) Percentage of animals with bacterial growth from the bone marrow. E) Bacterial load per gram bone marrow. F) Identification of cultured bacteria. CFU: Culture-forming units. n = 9-12 animals per group.

Next, we investigated bacterial adherence to small intestinal and colonic epithelium by fluorescence *in situ* hybridization (FISH) using both universal bacterial and *Pseudomonadota*-specific nucleotide probes [31]. Each specimen was carefully inspected under a confocal fluorescence microscope and assigned a semi-quantitative score by a blinded investigator based on abundance and location of fluorescent signal in the tissue. Among adhering bacteria, *Pseudomonadota* were as abundant as non-*Pseudomonadota* with no obvious differences between groups or intestinal segment (Suppl. Table S3). Irrespectively, we observed no statistical difference in general bacterial adherence (Fig. 3B-C, right panel).

Further, as a measure of bacterial translocation to the circulation, we aseptically collected femoral bone marrow biopsies, estimated the bacterial load by culturing and identified the colonies. The proportion of bone marrows colonized by bacteria and the mean estimated bacterial load appeared to be highest in the iFFT recipients, although not reaching statistical significance (CON-iFFT p=0.20, FFT-iFFT p=0.29) (Fig. 3D-E). Moreover, we identified five different bacterial species across bone marrows of iFFT recipients including streptococci and staphylococci, whereas FFT recipients were only colonized with *Enterococcus hirae*, an archetypical and highly abundant gut bacteria in preterm formula-fed piglets [20], [56] (Fig. 3F). Despite marginal differences, FFT restricted diverse bacterial translocation across the gut epithelium, an observation that was absent in the UV-inactivated virome.

### Inactivation of fecal filtrate preserves body weight gain and delays diarrhea onset

The growth of the animals was monitored by weighing shortly after birth and the following days, together with daily assessment of fecal and clinical status. We observed a decreased body weight gain in FFT recipients compared with CON and iFFT recipients as of study day four and five, which was also reflected in a marginally reduced average daily weight gain (p=0.05) (Fig. 4A-B). This was accompanied by an earlier onset of diarrhea in the FFT recipient group (mean 45 hours) relative to both the CON group (mean 67.7 hours) and iFFT recipient group (mean 65.3 hours) (Fig. 4C).

**Figure 4.**
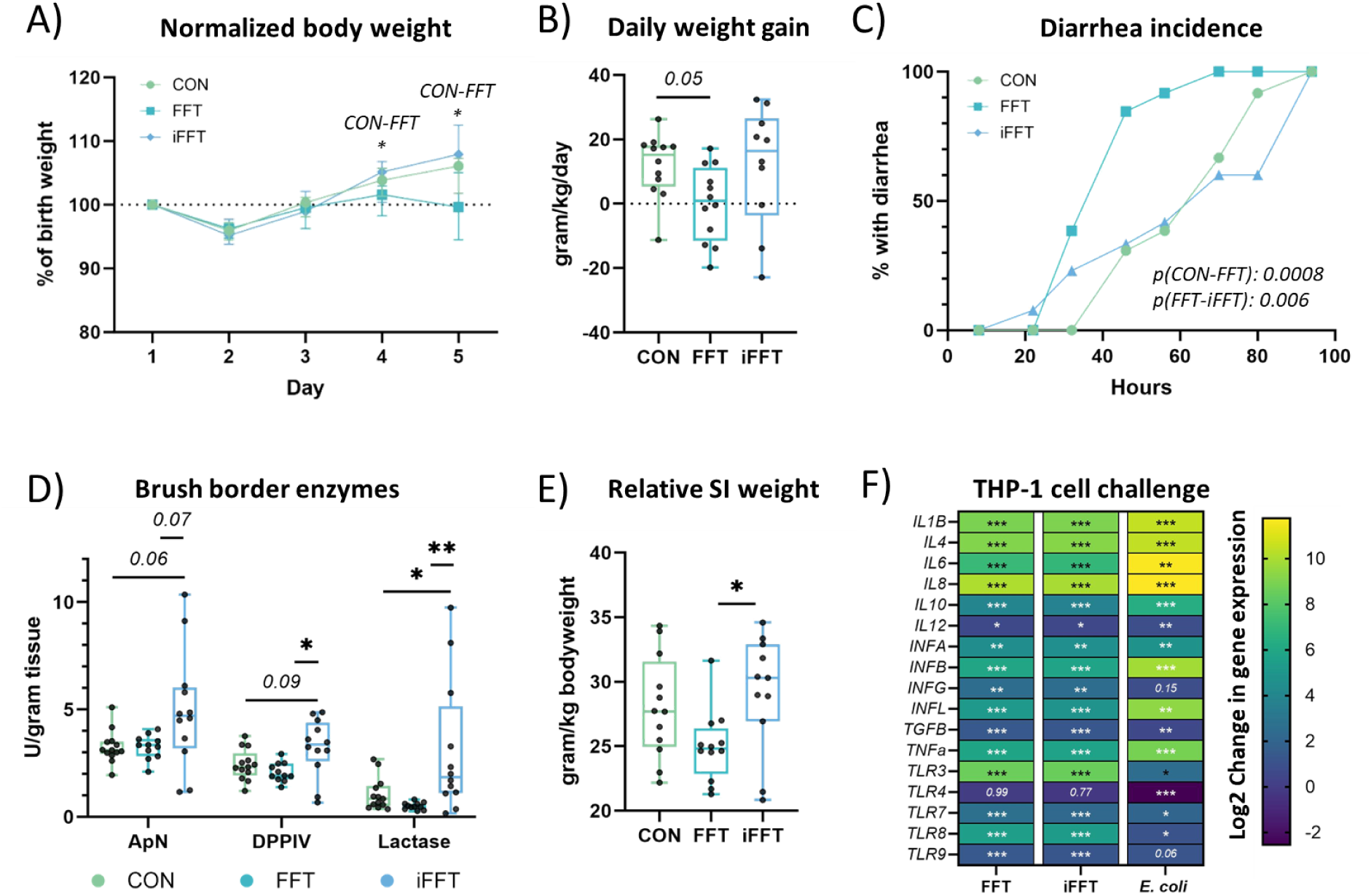
Preserved weight gain and delayed onset of diarrhea, but elevated brush border enzymes in recipients of inactivated fecal filtrate. A) Normalized body weight development over the study duration. B) Daily growth rate from birth to day five (gram per kg birth weight per day) C) Percentage of animals with diarrhea. D) Enzyme content in the small intestine tissue, ApN: Aminopeptidase N. DPPIV: Dipeptidyl peptidase IV. E) Small intestine weight relative to bodyweight at euthanasia. F) Normalized gene expression shown as log2(fold change) compared with unchallenged THP-1 cells. Data are mean values of three biological replicates. The continuous data is presented as boxplots with median and interquartile range. Significance levels are * p<0.05, ** p<0.01. n= 9-13 animals per group

Additionally, the activities of small intestinal brush-border enzymes aminopeptidase N, dipeptidyl peptidase-4 and lactase, assayed as a measure of mucosal function, were all elevated in iFFT recipients relative to both CON and FFT recipients, which were at similar levels (Fig. 4D, Suppl. table S4). This was in accordance with an increased small intestinal weight relative to bodyweight in iFFT recipients (Fig. 4E).

As induction of host immunity and side effects have previously been described following phage treatment [57], this prompted us to investigate if direct interaction between the donor fecal viruses and the mammalian immune system could underpin the observed differences between native FFT and iFFT. We challenged THP-1 derived macrophages with FFT, iFFT or sterile buffer and investigated the expression of several inflammatory genes standardized by the geometric average expression of reference genes *PGK1* and *ACTB.* Both the FFT and iFFT increased the gene expression of all assayed cytokines, as well as all nucleotide-binding toll-like receptors (TLRs 3, 7, 8 and 9) equally relative to unchallenged cells, suggesting that the UV-inactivation had no effect on the immunogenic potential (Fig. 4F). Likewise, our data indicated that the fecal filtrates were free of Gram-negative bacterial traces such as lipopolysaccharide, as no change in TLR4 expression was observed, contrary to a decrease observed in the cells challenged with a strain of *E. coli*. This strongly suggested that the side-effects observed after FFT were not due to immunogenic stimulation by viral epitopes.

### Transfer of native but not inactivated fecal filtrate constrains the gut bacteriome and increases virome heterogeneity in recipients

Having identified clearly distinct clinical responses in fecal filtrate recipients depending on the virome viability, we proceeded to investigate the impact on the recipient colonic bacteriome and virome by 16S rRNA gene amplicon and viral metagenomic sequencing, respectively. Although neither bacterial nor viral richness were affected by the FFT treatments or NEC status (Fig. 5A,D), we uncovered an interesting compositional pattern when inspecting the bacterial and viral ordination plots in combination (Fig. 5B,E). The native FFT with a viable virome constrained the resulting colonic bacterial composition into a smaller, homogeneous cluster, whereas the resulting virome composition of the same recipients turned more heterogeneous relative to control individuals (Fig. 5C,F). When the fecal filtrate was UV-inactivated prior to administration, it lost its constraining effect of the colonic bacteriome, and no concomitant diversification of the recipient virome was observed. We suggest delivery of viable phages via FFT, which modulate the bacterial composition in the recipient gut, as a potential model describing these trans-kingdom dynamics. However, the colonic microbiome composition of individuals receiving iFFT was not identical to controls either (Fig. 5B), suggesting that the UV irradiation might have had a collateral effect on the fecal filtrate with sufficient impact to influence microbiome composition.

**Figure 5.**
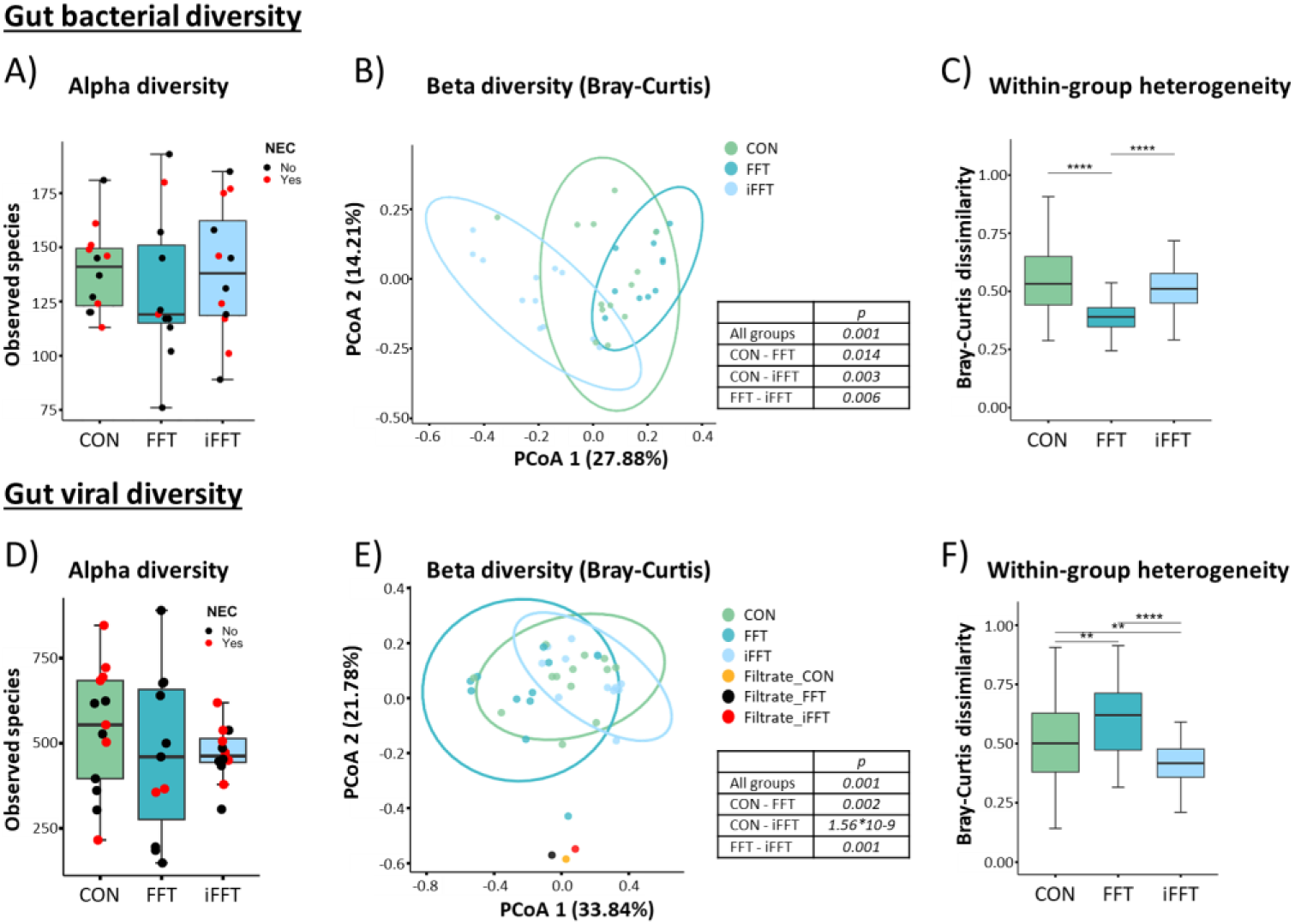
Active fecal filtrate changed the colonic bacterial composition and increased the virome heterogeneity. A) Colonic bacterial alpha diversity shown as number of observed species. B) Colonic bacterial composition as determined by Bray-Curtis dissimilarity metric. C) Colonic bacterial community heterogeneity. D) Colonic virome alpha diversity. E) Colonic virome composition as determined by Bray-Curtis dissimilarity metric. F) Colonic virome community heterogeneity. The data is presented as boxplots with median and interquartile range or Principal Coordinate Analysis (PCoA) ordination plots. Significance levels are ** p<0.01, *** p<0.0001, **** p<0.00001. n= 11-12 animals per group.

### Reduction of pathogenic bacteria by fecal filtrate transfer requires a viable virome

When inspecting bacterial composition on phylum level, the iFFT recipient group clearly stood out by having a significantly higher relative abundance of *Pseudomonadota*, to which numerous pathobionts belong, compared with FFT recipients (Fig. 6A). This difference was accompanied by a marked decrease in *Bacillota* (Suppl. Figure S1A, Suppl. Table S5). Further, when comparing differences in relative abundance among individual bacterial species, we observed a trend of reduced relative abundance of several NEC-related pathogens, e.g. *Clostridium perfringens* and *Escherichia* spp., in the FFT recipient group compared with CON recipient group (Fig. 6B, Suppl. Figure S1B, Suppl. Table S6). The iFFT recipients showed marked changes in the relative bacterial abundance, highlighted with increased relative abundance of pathobionts such as *Enterobacter hormaechi*, *Enterobacter* spp., *Clostridium perfringens*, *Klebsiella* spp., *Escherichia* compared with both CON and FFT recipients. Simultaneously, the change in microbiome showed reduced relative abundances of *Enterococcus faecium* and *Lactobacillus* spp. in the iFFT recipient group. With respect to the colonic virome, the FFT recipient group appeared markedly different from the remaining groups (Fig. 6C). Indeed, a comparison of the relative abundances of single viral OTUs revealed a clear enrichment of several viruses that were further classified as phages (unclassified Caudoviricetes and Siphoviridae) in the FFT recipient group relative to controls (Fig. 6D, Suppl. Table S7). Conversely, we observed a general reduction of phage relative abundance in the iFFT recipient group compared with FFT and CON recipients, of unclassified Caudovirales, Siphoviridae, Myoviridae, Autographiviridae, Microviridae and Cystoviridae. The relative abundance of Inoviridae and Podoviridae were further reduced in the iFFT compared with FFT recipient group. Host prediction revealed a high proportion of phages expected to infect *Enterococcae*, but in no clear pattern across the groups (Supplementary Figure 1C). Altogether, these data underline the demonstrated microbial diversity dynamics by showing colonic phage enrichment and reductions specifically in the abundance of NEC-associated bacteria only in individuals receiving native FFT. Conversely, the iFFT may still have perturbed the preterm colonic microbiome via unknown mechanisms.

**Figure 6.**
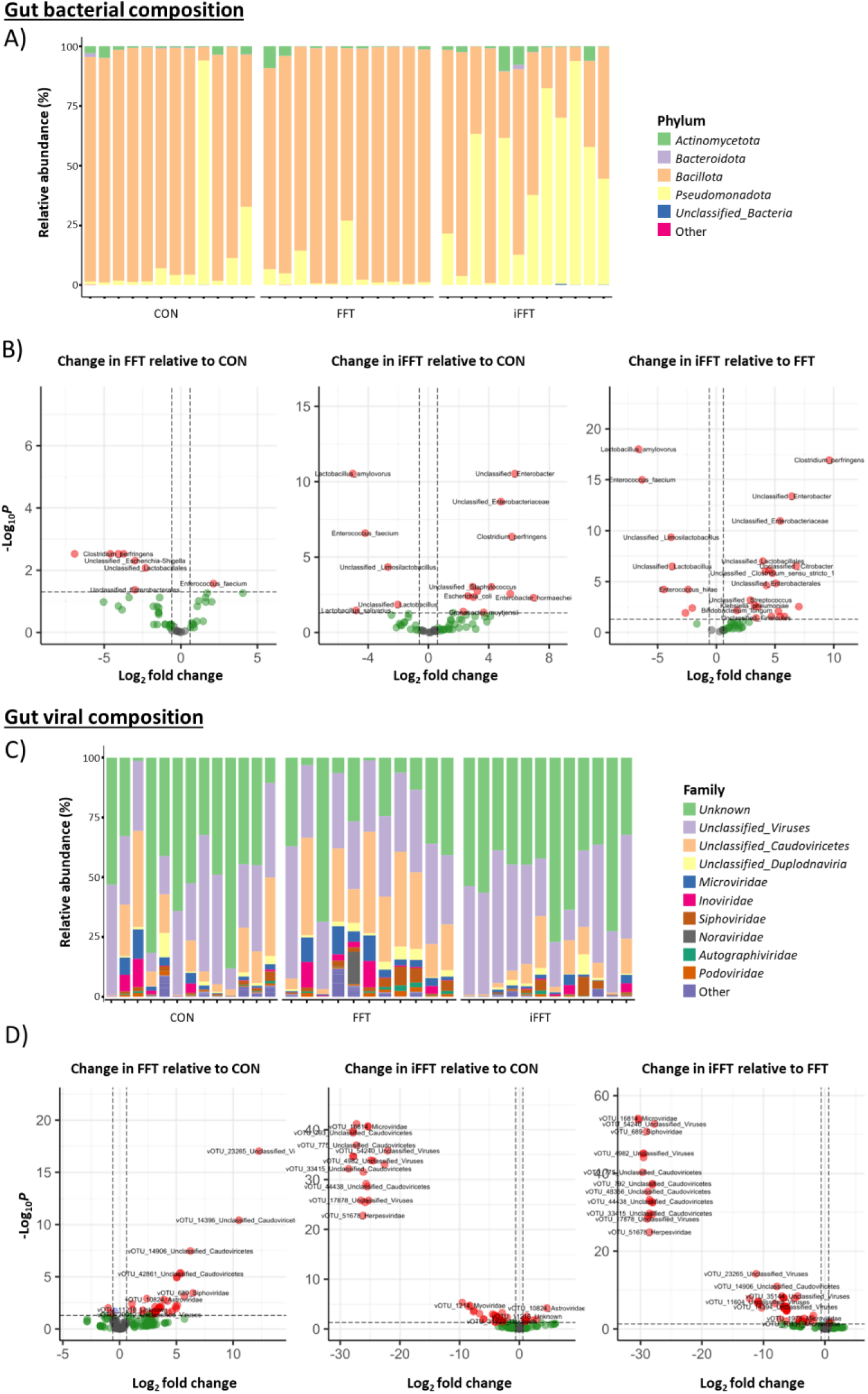
The active fecal filtrate reduced the relative abundances of NEC-associated bacteria, whereas the inactivated fecal filtrate promoted them. A) Colonic bacterial relative abundance. B) Most differently abundant colonic bacterial species. C) Colonic virome relative abundance and composition of recipients. D) Most differently abundant colonic virome signatures. Dots in red fulfill the requirements of significance level p<0.05 and log2(fold change)>0.6 in both directions. n= 11-12 animals per group.

## Discussion

Fecal filtrate transfer is suggested to work by introducing donor phages to restore recipient gut dysbiosis [58]. To date, no studies have investigated the importance of transferring phages in an active state by comparing the outcome with a UV-irradiated formulation. We demonstrate that UV-irradiation of a fecal filtrate effectively inactivates its virome and in turn abolishes the beneficial effects of FFT *in vivo*. When this was applied to a newborn piglet model of NEC, the native FFT constrained the gut microbiome to a core composition with fewer pathobionts and reduced colon pathology, while increasing the viral diversity in the recipients. Conversely, inactivated FFT did not constrain the gut microbiome and did not protect against NEC, but rather increased stomach lesion severity. Due to low acid secretion, the preterm piglet stomach is densely colonized with bacteria, reaching 10^8^ CFU per gram content [59]. This challenges the gastric mucosa to a degree where minor stimuli might initiate pathological processes. We speculate that photoinduced damage to UV-irradiated fecal filtrate molecules beyond viral DNA might have created detrimental reactive species. However, this was not obvious from the immune cell stimulations *in vitro,* which showed similar immune responses towards native and inactivated fecal filtrate.

Meanwhile, native FFT also induced side effects that manifested as early onset of diarrhea and reduced weight gain, whereas these were not observed in recipients of inactivated FFT. Hence, both NEC protection and side effects appeared to be associated with an active transferred virome. As both filtrates stimulated immune cells in a similar pattern, this suggests that fecal filtrate acts *in vivo* by modulating the gut microbiome through the actions of an active virome, rather than directly stimulating immune cells.

The presence of phages in the gut might provide an additional layer of protection from invasive bacterial pathogens. Barr and colleagues described that phages adhering to mucus actively protect the epithelium from bacterial invasion and observed that this effect was dependent on active phage infection and replication [60]. We observed changes in the colonic microbiome on bacterial and viral level in FFT recipients, specifically with decrease in *Enterobacteriaceae, Klebsiella spp.* and *Clostridioides perfringens*. Oppositely, these bacterial signatures were increased in iFFT recipients, along with a marked increase in *Pseudomonadota*. The shift in bacterial composition preceding NEC has been extensively described, showing reduced diversity with increased relative abundance of *Pseudomonadota*, and coupled with decrease in obligate anaerobes such as *Bacteroidota* [4], [8], [11], [12], [13], [61], [62]. [63] On a species level, increased abundance of *C. perfringens* and *K. pneumoniae* are evident prior to NEC diagnosis [64]. The overall bacterial diversity is a main factor in the dysbiosis preceding NEC, with complete dominance of either *Bacillota* or *Pseudomonadota* [65] or specific opportunistic pathogenic species such as *E. coli* or *Enterobacter* [7]. This underlines the importance of not only *Pseudomonadota* but also overall bacterial diversity.

In this study, UV-inactivated fecal filtrate was ineffective in protecting against NEC. The loss in efficiency following virus inactivation emphasizes the importance of using fresh donor material, and that attention should be paid to limiting the damage to the viral particles. This does not seem as essential for the efficacy of FMT, where comparable effects for NEC protection and reduced inflammation were observed for both fresh FMT and UV-irradiated FMT [66], suggesting that the effect might be mediated through the bacterial epitopes, contrary to FFT.

The strengths of this study lie in the use of our preterm piglet model which resembles preterm infants developmentally and clinically, especially in the context of NEC [67]. Even so, we are aware of several limitations; The FFT was inactivated by UV-irradiation, and we determined the duration based on the inactivation of a known mock phage community. Total phage inactivation was observed at 30 minutes exposure, and to ensure complete viral inactivation of the fecal filtrate solution, we decided to increase the time to 45 minutes. This increase might have interfered with bioactive molecules in the fecal filtrate, for example volatile organic compounds (VOC), a group of diverse compounds with broad effects in humans and bacteria [68], that might be involved in intestinal diseases such as NEC [69], [70]. Albeit, UV inactivation is a more focused and gentle viral inactivation procedure relative to heat treatment, and we confirmed the presence of viral DNA with fluorescence microscopy, suggesting that the viral capsids remained intact after UV-irradiation. However, for future investigations we suggest to instead inactivate a purified fecal virus fraction devoid of smaller molecules to prevent the generation of photo-induced reactive species potentially interfering with the clinical endpoints [28].

We expected the iFFT recipients to resemble the CON recipients, but while the clinical phenotype was similar between the two groups, the iFFT recipients displayed a diverged bacteriome and virome composition. Prior studies transferring a heat-inactivated virome in mice observed no change in microbiome composition of recipients [71], and have supported the argument for the importance of live phages for modulation of the microbiome. In another study, a heat-and nuclease treated fecal filtrate was administered as control against an active filtrate in a murine model of virome-based recovery of antibiotic-induced dysbiosis [72]. In the cell assay, we used monocytes although enterocytes would have been more clinically relevant, especially in relation to the diarrheal side effects. As our aim was to investigate the immune response towards the fecal filtrates and not mimic the gut environment, we chose the monocytes based on their phagocytic capacity of the cells and range of inflammatory genes produced.

In summary, we have shown that FFT efficacy depends on active donor phages to provide protection against NEC, potentially due to engrafting donor phages interacting with the resident gut microbiome, and unlikely due to direct phage-host interplay.

## Acknowledgements

The THP-1 cell line was supplied by Professor John Elmerdahl Olsen, section for Veterinary Clinical Microbiology, University of Copenhagen. The fluorescence microscopy was performed at the Core Facility for Integrated Microscopy, Faculty of Health and Medical Sciences, University of Copenhagen.

## Data availability

Data from the 16S rRNA gene sequencing and metavirome sequencing were submitted to the BioProject database with the accession number PRJNA1032308, and are available at: https://www.ncbi.nlm.nih.gov/bioproject/PRJNA1032308/

