## Supplementary material for "Protection against necrotizing enterocolitis by fecal filtrate transfer requires an active donor virome"

### Supplementary Figures

#### Supplementary Figure S1:

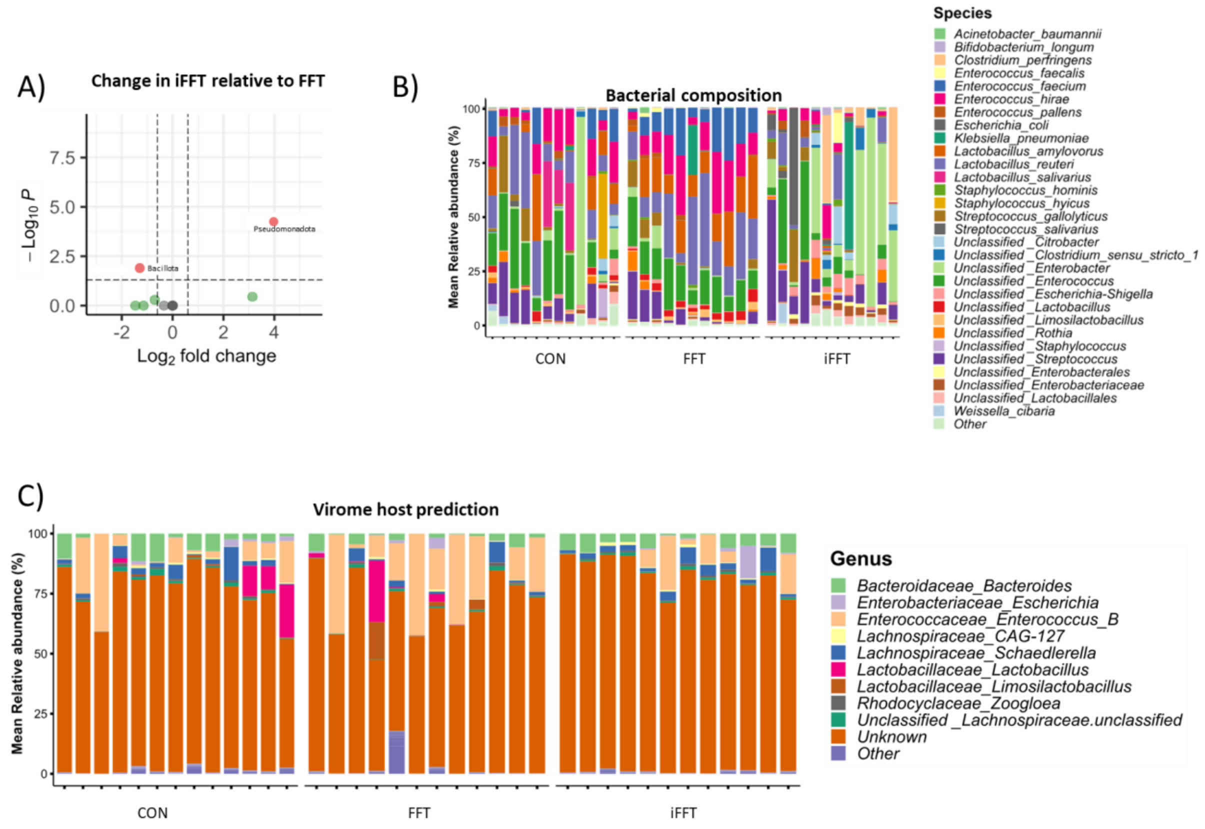


**Supplementary figure S1.** Administration of inactivated FFT changes the microbiome from FFT. A) Phylum-level changes in colon microbiome of iFFT recipients compared with FFT. Dots in red fulfill the requirements of significance level p<0.05 and log2(fold change)>0.6 in both directions. B) Colon bacterial relative abundance in recipients. C) Host prediction of colon virome in recipients.

### Supplementary Tables

#### Supplementary table S1: UV-inactivation of mock bacteriophages

| | | **Phage titer (PFU/ml) after UV exposure** | | | |
|---|---|---|---|---|---|
| **Phage** | **Host strain (reference)** | *0 min* | *10min* | *20min* | *30min* |
| ϕ29 | *Bacillus subtilis* (DSM 5547) | $1.10 \times 10^6$ | 0 | 0 | 0 |
| T4 | *E.coli* (DSM 613) | $1.50 \times 10^5$ | 0 | 0 | 0 |
| c2 | *Lactococcus lactis* (MG 1363) | $3.30 \times 10^6$ | 0 | 20 | 0 |
| ϕ6 | *Pseudomonas syringae* (DSM 21482) | $1.5 \times 10^3$ | 0 | 0 | 0 |
| ϕ2972 | *S. thermophiles* (DGCC 7710) | $1.21 \times 10^6$ | 50 | 0 | 0 |
| ϕX174 | *E. coli* (ATTC 13706) | $8.70 \times 10^5$ | 0 | 0 | 0 |

*PFU/ml: Plaque-forming units per ml*

#### Supplementary table S2: Primer sequences

| Gene target | Forward primer | Reverse primer | Reference |
|---|---|---|---|
| *IL1B* | GGCCACATTTGGTTCTAAGAAA | TAAATAGGGAAGCGGTTGCTC | (Van Belleghem et al., 2017) |
| *IL4* | GCCCTAAACAGATGAAGTGCTC | CCATGGCCACAACAACTGAC | Designed from Primer-BLAST (Ye et al., 2012) |
| *IL6* | GGTACATCCTCGACGGCATC | GCCTCTTTGCTGCTTTCACAC | (Van Belleghem et al., 2017) |
| *IL8* | AGTTTTTGAAGAGGGCTGAGAAT | GCTTGAAGTTTCACTGGCATCT | (Kawarizadeh et al., 2021) |
| *IL10* | CAACCTGCCTAACATGCTTCGAGAT | CCTCCAGCAAGGACTCCTTTAACAAC | (Y. Li et al., 2022) |
| *IL12* | GCGGAGCTGCTACACTCTCT | GGTGGGTCAGGTTTGATGAT | (Zhang et al., 2016) |
| *INFA* | ATTTCTGCTCTGACAACCTC | TGACAGAGACTCCCCTGATG | (Chen et al., 2012) |
| *INFB* | AAACTCATGAGCAGTCTGCA | AGGAGATCTTCAGTTTCGGAGG | (Richtsteiger et al., 2003) |
| *INFG* | GACCAGAGCATCCAAAAGAGT | ATTGCTTTGCGTTGGACATTC | (Meenakshi et al., 2016) |
| *INFL* | GGACGCCTTGGAAGAGTCACT | AGAAGCCTCAGGTCCCAATTC | (Slater et al., 2010) |
| *TGFB* | GAAGGGAGACAATCGCTTTAGC | TGTAGACTCCTTCCCGGTTGAG | (Van Belleghem et al., 2017) |
| *TNFA* | CCCAGGGACCTCTCTCTAATC | ATGGGCTACAGGCTTGTCACT | (Van Belleghem et al., 2017) |
| *TLR3* | AACTCCATCTCATGTCCAACTCAA | GATGACAAGCCATTATGAGACAGATC | (Månsson et al., 2010) |
| *TLR4* | CAGAGTTGCTTTCAATGGCATC | AGACTGTAATCAAGAACCTGGAGG | (Tedesco et al., 2018) |
| *TLR7* | GGAGGTATTCCCACGAACACC | TGACCCCAGTGGAATAGGTACAC | (Vrećа et al., 2018) |
| *TLR8* | AACTTTCTATGATGCTTACATTTCTTATGAC | GGTGGTAGCGCAGCTCATTT | (B. Li et al., 2013) |
| *TLR9* | GTCCAGCACTCGGAGGTTTC | TGGTGTTGAAGGACAGTTCTCTCT | (Mortezagholi et al., 2017) |
| *ACTB* | AAGGTGACAGCAGTCGGTT | TGTGTGGACTTGGGAGAGG | (Wei et al., 2023) |
| *PGK1* | CAAGAAGTATGCTGAGGCTGTCA | CAAATACCCCCACAGGACCAT | (Falkenberg et al., 2011) |

#### Supplementary table S3: Bacterial types observed by fluorescence microscopy

| | Small intestine | Colon |
|---|---|---|

| | *CON* | *FFT* | *iFFT* | *CON* | *FFT* | *iFFT* |
|---|---|---|---|---|---|---|
| All bacteria, rods | 69.2 | 66.7 | 75 | 76.9 | 83.3 | 100 |
| All bacteria, cocci | 69.2 | 66.7 | 75 | 23.1 | 50 | 41.6 |
| *Pseudomonadota* | 76.9 | 66.7 | 83.3 | 76.9 | 66.7 | 83.3 |

*% of tissues where the type was observed. n=12-13 in each group*

#### Supplementary table S4: Intestinal brush border enzyme activity

| | **CON** | **FFT** | **iFFT** |
|---|---|---|---|
| ApA | 3.25 ± 0.75 | 3.25 ± 0.56 | 4.93 ± 2.73 |
| ApN | 1.42 ± 0.31 | 1.56 ± 0.33 | 2.01 ± 1.24 |
| DPPIV | 2.43 ± 0.71 | 2.07 ± 0.44 | 3.23 ± 1.35 |
| Sucrase | 0.060 ± 0.026 | 0.051 ± 0.013 | 0.051 ± 0.033 |
| Maltase | 1.01 ± 0.41 | 1.14 ± 0.32 | 0.98 ± 0.47 |
| Lactase | 1.02 ± 0.73 | 0.51 ± 0.14 | 2.89 ± 2.85 |

*Enzyme activity levels in U/gram of tissue. Mean ± SD.*
*ApA: Aminopeptidase A. ApN: Aminopeptidase N. DPPIV: Dipeptidyl peptidase IV. n=11-13 in each group*

#### Supplementary table S5: Change in 16S phylum level in iFFT recipients relative to FFT recipients

| **Phylum** | **Fold change** | **Log2(fold change)** | **Adjusted p-value** |
|---|---|---|---|
| Firmicutes | 0.41 | -1.28 | *0.013* |
| Proteobacteria | 15.67 | 3.97 | *0.00006* |

#### Supplementary table S6: Change in 16S genus level

##### Change in FFT relative to CON

| **Phylum** | **Species** | **Fold change** | **Log2(fold change)** | **Adjusted p-value** |
|---|---|---|---|---|
| Firmicutes | Enterococcus_faecalis | 16.55931 | 4.049571 | *0.053929* |
| Firmicutes | Enterococcus_faecium | 4.374098 | 2.128986 | *0.026737* |
| Firmicutes | Unclassified _Lactobacillus | 3.309794 | 1.726741 | *0.053929* |
| Firmicutes | Lactobacillus_amylovorus | 3.051587 | 1.60956 | *0.074808* |
| Firmicutes | Unclassified _Limosilactobacillus | 2.252684 | 1.171645 | *0.096943* |
| Actinobacteriota | Bifidobacterium_longum | 0.38523 | -1.37621 | *0.074808* |
| Firmicutes | Unclassified_Lactobacillales | 0.202912 | -2.30107 | *0.008421* |
| Proteobacteria | Unclassified _Escherichia-Shigella | 0.126363 | -2.98435 | *0.004943* |
| Proteobacteria | Unclassified_Enterobacterales | 0.126167 | -2.98659 | *0.04257* |
| Firmicutes | Clostridioides_difficile | 0.096378 | -3.37516 | *0.074808* |
| Proteobacteria | Klebsiella_pneumoniae | 0.075412 | -3.72907 | *0.00298* |
| Firmicutes | Clostridium_perfringens | 0.059086 | -4.08103 | *0.00298* |

| | | | | |
|---|---|---|---|---|
| Proteobacteria | Unclassified _Citrobacter | 0.041497 | -4.59086 | *0.00298* |
| Firmicutes | Lactobacillus_salivarius | 0.00821 | -6.92842 | *0.00298* |

#### Change in iFFT relative to CON

| Phylum | Species | Fold change | Log2(fold change) | Adjusted p-value |
|---|---|---|---|---|
| Proteobacteria | Enterobacter_hormaechei | 124.8599 | 6.964166 | *0.00492* |
| Proteobacteria | Unclassified _Enterobacter | 53.09797 | 5.730585 | *2.91E-11* |
| Firmicutes | Clostridium_perfringens | 46.19957 | 5.529807 | *4.37E-07* |
| Firmicutes | Enterococcus_faecalis | 42.78497 | 5.419032 | *0.002674* |
| Proteobacteria | Unclassified_Enterobacteriaceae | 27.91378 | 4.802906 | *2.09E-09* |
| Firmicutes | Streptococcus_salivarius | 18.04417 | 4.173461 | *0.000941* |
| Proteobacteria | Unclassified _Klebsiella | 17.28909 | 4.11179 | *0.084761* |
| Proteobacteria | Unclassified _Moraxella | 13.48229 | 3.752994 | *0.091316* |
| Proteobacteria | Cronobacter_muytjensii | 12.65627 | 3.66178 | *0.046798* |
| Firmicutes | Unclassified _Blautia | 9.488757 | 3.246219 | *0.001729* |
| Firmicutes | Unclassified_Firmicutes | 8.53791 | 3.093883 | *0.08678* |
| Firmicutes | Unclassified _Clostridium_sensu_stricto_1 | 7.899455 | 2.981753 | *0.004496* |
| Firmicutes | Unclassified _Staphylococcus | 7.606938 | 2.927316 | *0.000927* |
| Proteobacteria | Escherichia_coli | 6.132408 | 2.616454 | *0.003806* |
| Proteobacteria | Unclassified _Acinetobacter | 5.671873 | 2.503825 | *0.069843* |
| Firmicutes | Staphylococcus_hominis | 4.935646 | 2.303239 | *0.041168* |
| Firmicutes | Unclassified_Lactobacillales | 3.144761 | 1.65295 | *0.050265* |
| Firmicutes | Enterococcus_hirae | 0.400766 | -1.31917 | *0.058363* |
| Firmicutes | Lactobacillus_reuteri | 0.396259 | -1.33549 | *0.098768* |
| Firmicutes | Lactobacillus_crispatus | 0.182612 | -2.45314 | *0.069843* |
| Firmicutes | Unclassified _Limosilactobacillus | 0.155788 | -2.68234 | *4.6E-05* |
| Firmicutes | Enterococcus_faecium | 0.054523 | -4.197 | *2.58E-07* |
| Firmicutes | Lactobacillus_salivarius | 0.036094 | -4.79211 | *0.034089* |
| Firmicutes | Lactobacillus_amylovorus | 0.030978 | -5.01262 | *2.91E-11* |

#### Change in iFFT relative to FFT

| Phylum | Species | Fold change | Log2(fold change) | Adjusted p-value |
|---|---|---|---|---|
| Firmicutes | Clostridium_perfringens | 781.8975 | 9.610836 | *1.16E-17* |
| Proteobacteria | Enterobacter_hormaechei | 130.3412 | 7.026149 | *0.002869* |
| Proteobacteria | Unclassified _Citrobacter | 111.3841 | 6.7994 | *3.06E-07* |
| Proteobacteria | Unclassified _Enterobacter | 85.74937 | 6.422054 | *4.11E-14* |
| Unclassified_Bacteria | Unclassified_Bacteria | 55.79616 | 5.802094 | *0.02678* |
| Proteobacteria | Unclassified_Enterobacteriaceae | 43.16986 | 5.431952 | *1.09E-11* |
| Proteobacteria | Unclassified _Kosakonia | 42.77008 | 5.41853 | *0.02678* |
| Proteobacteria | Unclassified _Moraxella | 39.16081 | 5.291339 | *0.008225* |
| Proteobacteria | Unclassified_Enterobacterales | 33.62919 | 5.071642 | *1.5E-05* |

| | | | | |
|---|---|---|---|---|
| Firmicutes | Unclassified _Clostridium_sensu_stricto_1 | 27.32367 | 4.772079 | *1.48E-06* |
| Proteobacteria | Unclassified _Klebsiella | 25.7615 | 4.687145 | *0.02678* |
| Proteobacteria | Unclassified _Escherichia-Shigella | 22.5529 | 4.495241 | *7.25E-07* |
| Firmicutes | Unclassified _Blautia | 19.23727 | 4.265832 | *2.39E-05* |
| Proteobacteria | Escherichia_coli | 17.86092 | 4.158734 | *9.77E-07* |
| Firmicutes | Unclassified_Lactobacillales | 15.49814 | 3.954023 | *9.7E-08* |
| Firmicutes | Streptococcus_salivarius | 11.4997 | 3.523525 | *0.002897* |
| Firmicutes | Unclassified_Firmicutes | 10.42614 | 3.382134 | *0.036214* |
| Proteobacteria | Klebsiella_pneumoniae | 9.518581 | 3.250746 | *0.002252* |
| Firmicutes | Unclassified _Streptococcus | 7.444754 | 2.896224 | *0.000648* |
| Proteobacteria | Unclassified _Acinetobacter | 4.717036 | 2.237881 | *0.075851* |
| Actinobacteriota | Bifidobacterium_longum | 3.346247 | 1.742544 | *0.006781* |
| Firmicutes | Lactobacillus_reuteri | 0.241506 | -2.04987 | *0.004007* |
| Firmicutes | Enterococcus_hirae | 0.189402 | -2.40048 | *5.99E-05* |
| Firmicutes | Unclassified_Lactobacillaceae | 0.161076 | -2.63419 | *0.011858* |
| Firmicutes | Unclassified _Lactobacillus | 0.073881 | -3.75865 | *3.2E-07* |
| Firmicutes | Unclassified _Limosilactobacillus | 0.069157 | -3.85399 | *4.48E-10* |
| Firmicutes | Lactobacillus_crispatus | 0.045179 | -4.46821 | *5.99E-05* |
| Firmicutes | Enterococcus_faecium | 0.012465 | -6.32598 | *1.01E-15* |
| Firmicutes | Lactobacillus_amylovorus | 0.010151 | -6.62218 | *9.87E-19* |

Supplementary table S7: Change in virome

#### Change in FFT relative to CON

| Phylum | Species | Fold change | Log2(fold change) | Adjusted p-value |
|---|---|---|---|---|
| Pisuviricota | Astroviridae_Porcine_astrovirus_3 | 4925.52 | 12.26606 | *1.36E-15* |
| Uroviricota | Unclassified_Caudoviricetes | 1451.27 | 10.5031 | *2.8E-09* |
| Uroviricota | Unclassified_Caudoviricetes | 88.19 | 6.462632 | *0.00647* |
| Unclassified_Viruses | Unclassified_Viruses | 75.06 | 6.230062 | *1.68E-06* |
| Uroviricota | Unclassified_Caudoviricetes | 52.30 | 5.708831 | *0.01135* |
| Unclassified_Viruses | Unclassified_Viruses | 41.73 | 5.382935 | *0.00015* |
| Uroviricota | Unclassified_Caudoviricetes | 40.66 | 5.345627 | *0.00015* |
| Unknown | Unknown | 32.92 | 5.040805 | *0.00024* |
| Unclassified_Viruses | Unclassified_Viruses | 32.62 | 5.027523 | *0.00025* |
| Unclassified_Viruses | Unclassified_Viruses | 18.51 | 4.210244 | *0.0261* |
| Uroviricota | Unclassified _Siphoviridae | 14.77 | 3.884955 | *0.02021* |
| Hofneiviricota | Inoviridae_environmental samples | 5.23 | 2.387168 | *0.01830* |
| Unclassified_Viruses | Unclassified_Viruses | 2.11 | 1.078271 | *0.04260* |

#### Change in iFFT relative to CON

| Phylum | Species | Fold change | Log2(fold change) | Adjusted p-value |
|---|---|---|---|---|
| Pisuviricota | Astroviridae_Porcine_astrovirus_3 | 24.65 | 4.623371 | *0.000473* |
| Unclassified_Viruses | Unclassified_Viruses | 7.45 | 2.897503 | *0.043437* |
| Unknown | Unknown | 2.40 | 1.265094 | *0.015889* |
| Unknown | Unknown | 1.83 | 0.874816 | *0.04227* |
| Unclassified_Viruses | Unclassified_Viruses | 0.24 | -2.04084 | *0.02249* |
| Uroviricota | Unclassified _Siphoviridae | 0.14 | -2.87025 | *0.00349* |
| Unclassified_Viruses | Unclassified_Viruses | 0.10 | -3.27089 | *0.046071* |
| Unclassified_Viruses | Unclassified_Viruses | 0.10 | -3.31114 | *0.046071* |
| Unclassified_Viruses | Unclassified_Viruses | 0.10 | -3.3326 | *0.049679* |
| Unclassified_Viruses | Unclassified_Viruses | 0.09 | -3.4615 | *0.033221* |
| Uroviricota | Unclassified_Caudoviricetes | 0.09 | -3.47638 | *0.046071* |
| Uroviricota | Unclassified _Autographiviridae | 0.09 | -3.52755 | *0.04227* |
| Unclassified_Viruses | Unclassified_Viruses | 0.08 | -3.69166 | *0.016889* |
| Uroviricota | Unclassified_Caudoviricetes | 0.07 | -3.76072 | *0.019174* |
| Unclassified_Viruses | Unclassified_Viruses | 0.07 | -3.76946 | *0.022797* |
| Unclassified_Duplodnaviria | Unclassified_Duplodnaviria | 0.07 | -3.84658 | *0.028147* |
| Uroviricota | Unclassified _Siphoviridae | 0.06 | -3.99139 | *0.009694* |
| Unclassified_Viruses | Unclassified_Viruses | 0.05 | -4.23892 | *0.009694* |
| Uroviricota | Unclassified _Podoviridae | 0.05 | -4.46636 | *0.006316* |
| Unclassified_Duplodnaviria | Unclassified_Duplodnaviria | 0.02 | -5.95126 | *0.043117* |
| Uroviricota | Unclassified _Amalieviridae | 0.02 | -5.96453 | *0.043437* |
| Unclassified_Viruses | Unclassified_Viruses | 0.01 | -6.65769 | *0.005112* |
| Uroviricota | Unclassified_Caudoviricetes | 0.005 | -7.60297 | *0.001258* |
| Uroviricota | Unclassified_Caudoviricetes | 0.005 | -7.65755 | *0.002218* |
| Uroviricota | Unclassified _Myoviridae | 0.003 | -8.26818 | *0.000152* |
| Phixviricota | Unclassified _Microviridae | 0.002 | -8.48768 | *0.000201* |
| Uroviricota | Unclassified_Caudoviricetes | 0.001 | -9.54816 | *4.1E-05* |
| Unclassified_Viruses | Unclassified_Viruses | 2.3E-07 | -22.0492 | *3.82E-35* |
| Uroviricota | Unclassified _Siphoviridae | 1.62E-07 | -22.5589 | *1.28E-32* |
| Unclassified_Viruses | Unclassified_Viruses | 4.29E-08 | -24.474 | *2.54E-33* |
| Uroviricota | Unclassified_Caudoviricetes | 3.3E-08 | -24.8532 | *2.11E-33* |
| Unclassified_Viruses | Unclassified_Viruses | 2.52E-08 | -25.2436 | *1.37E-25* |
| Phixviricota | Unclassified _Microviridae | 2.39E-08 | -25.3196 | *1.27E-39* |
| Unclassified_Viruses | Unclassified_Viruses | 2.28E-08 | -25.3845 | *1.27E-39* |
| Uroviricota | Unclassified_Caudoviricetes | 2.13E-08 | -25.4826 | *2.64E-28* |
| Duplornaviricota | Cystoviridae_Cystovirus | 1.9E-08 | -25.6519 | *7.78E-29* |
| Unclassified_Viruses | Unclassified_Viruses | 1.69E-08 | -25.8151 | *3.23E-25* |
| Uroviricota | Unclassified_Caudoviricetes | 1.37E-08 | -26.12 | *3.46E-31* |

| | | | | |
|---|---|---|---|---|
| Peploviricota | Unclassified _Herpesviridae | 1.33E-08 | -26.1626 | *1.41E-22* |
| Unclassified_Viruses | Unclassified_Viruses | 1.04E-08 | -26.5247 | *1.37E-25* |
| Unclassified_Viruses | Unclassified_Viruses | 6.21E-09 | -27.263 | *1.04E-39* |
| Uroviricota | Unclassified_Caudoviricetes | 6.13E-09 | -27.2819 | *3.39E-36* |
| Uroviricota | Unclassified_Caudoviricetes | 4.2E-09 | -27.8273 | *4.18E-34* |
| Unclassified_Viruses | Unclassified_Viruses | 4.19E-09 | -27.8288 | *4.18E-34* |
| Uroviricota | Unclassified_Caudoviricetes | 4.15E-09 | -27.8454 | *1.29E-38* |
| Uroviricota | Unclassified_Caudoviricetes | 4.08E-09 | -27.8679 | *7.25E-39* |
| Uroviricota | Unclassified_Caudoviricetes | 2.48E-09 | -28.5858 | *8.64E-32* |

#### Change in iFFT relative to FFT

| Phylum | Species | Fold change | Log2(fold change) | Adjusted p-value |
|---|---|---|---|---|
| Uroviricota | Unclassified_Caudoviricetes | 1.93 | 0.950935 | *0.033643* |
| Unclassified_Viruses | Unclassified_Viruses | 0.27 | -1.86737 | *0.034622* |
| Uroviricota | Unclassified _Siphoviridae | 0.27 | -1.89441 | *0.000979* |
| Uroviricota | Unclassified _Myoviridae | 0.22 | -2.18038 | *0.005775* |
| Unclassified_Viruses | Unclassified_Viruses | 0.12 | -3.02151 | *0.038343* |
| Unclassified_Viruses | Unclassified_Viruses | 0.09 | -3.37763 | *0.005281* |
| Uroviricota | Unclassified_Caudoviricetes | 0.09 | -3.40842 | *0.033643* |
| Unclassified_Viruses | Unclassified_Viruses | 0.09 | -3.45427 | *0.03345* |
| Uroviricota | Unclassified_Caudoviricetes | 0.08 | -3.52964 | *0.032455* |
| Uroviricota | Unclassified_Caudoviricetes | 0.07 | -3.79662 | *0.009996* |
| Uroviricota | Unclassified_Caudoviricetes | 0.06 | -4.13779 | *0.046503* |
| Phixviricota | Unclassified _Microviridae | 0.05 | -4.29759 | *0.008359* |
| Unclassified_Viruses | Unclassified_Viruses | 0.05 | -4.42801 | *2.66E-08* |
| Phixviricota | Unclassified _Microviridae | 0.03 | -4.9211 | *8.69E-08* |
| Unknown | Unknown | 0.01 | -6.14105 | *3.93E-05* |
| Uroviricota | Unclassified _Podoviridae | 0.01 | -6.1989 | *3.37E-05* |
| Unclassified_Viruses | Unclassified_Viruses | 0.01 | -6.2269 | *2.38E-05* |
| Uroviricota | Unclassified_Caudoviricetes | 0.01 | -6.22854 | *1.94E-05* |
| Uroviricota | Unclassified_Caudoviricetes | 0.01 | -6.22992 | *3.6E-05* |
| Unclassified_Viruses | Unclassified_Viruses | 0.01 | -6.24091 | *6.53E-05* |
| Unclassified_Viruses | Unclassified_Viruses | 0.01 | -6.26229 | *6.8E-05* |
| Unclassified_Viruses | Unclassified_Viruses | 0.012852606 | -6.2818 | *1.51E-05* |
| Unclassified_Viruses | Unclassified_Viruses | 0.012749906 | -6.29337 | *6.92E-06* |
| Uroviricota | Unclassified _Siphoviridae | 0.012601385 | -6.31027 | *0.024063* |
| Unknown | Unknown | 0.012332017 | -6.34145 | *7.95E-06* |
| Unclassified_Viruses | Unclassified_Viruses | 0.012307621 | -6.3443 | *1.13E-05* |
| Unclassified_Viruses | Unclassified_Viruses | 0.012167653 | -6.36081 | *1.52E-05* |
| Uroviricota | Unclassified_Caudoviricetes | 0.012058393 | -6.37382 | *1.71E-05* |
| Unclassified_Viruses | Unclassified_Viruses | 0.012043923 | -6.37555 | *1.14E-05* |

| | | | | |
|---|---|---|---|---|
| Uroviricota | Unclassified_Caudoviricetes | 0.011826564 | -6.40183 | *8.6E-06* |
| Uroviricota | Unclassified_Caudoviricetes | 0.011692539 | -6.41827 | *1.51E-05* |
| Unclassified_Viruses | Unclassified_Viruses | 0.011244908 | -6.47458 | *7.11E-06* |
| Uroviricota | Unclassified _Autographiviridae | 0.011069062 | -6.49732 | *1.13E-05* |
| Uroviricota | Unclassified_Caudoviricetes | 0.010849779 | -6.52619 | *1.13E-05* |
| Unclassified_Duplodnaviria | Unclassified_Duplodnaviria | 0.010286978 | -6.60304 | *6.3E-06* |
| Unknown | Unknown | 0.009924475 | -6.65479 | *1.13E-05* |
| Uroviricota | Unclassified_Caudoviricetes | 0.009752216 | -6.68005 | *6E-08* |
| Uroviricota | Unclassified_Caudoviricetes | 0.009745406 | -6.68106 | *7.38E-08* |
| Uroviricota | Unclassified_Caudoviricetes | 0.009592216 | -6.70392 | *1.36E-05* |
| Unknown | Unknown | 0.009586605 | -6.70476 | *1.52E-05* |
| Uroviricota | Unclassified _Siphoviridae | 0.008824685 | -6.82424 | *6.41E-06* |
| Uroviricota | Unclassified_Caudoviricetes | 0.008791941 | -6.8296 | *8.49E-08* |
| Uroviricota | Unclassified_Caudoviricetes | 0.007863002 | -6.9907 | *2.45E-08* |
| Uroviricota | Unclassified_Caudoviricetes | 0.006490849 | -7.26738 | *0.008047* |
| Uroviricota | Unclassified _Siphoviridae | 0.004455343 | -7.81025 | *8.41E-11* |
| Unclassified_Viruses | Unclassified_Viruses | 0.003647237 | -8.09898 | *1.44E-06* |
| Uroviricota | Unclassified_Caudoviricetes | 0.000794415 | -10.2978 | *1.29E-05* |
| Phixviricota | Unclassified _Microviridae | 0.000590661 | -10.7254 | *1.85E-06* |
| Peploviricota | Unclassified _Herpesviridae | 0.00055472 | -10.816 | *4.19E-07* |
| Unclassified_Viruses | Unclassified_Viruses | 0.000537191 | -10.8623 | *1.23E-06* |
| Unclassified_Viruses | Unclassified_Viruses | 0.000503724 | -10.9551 | *6.22E-07* |
| Unclassified_Viruses | Unclassified_Viruses | 0.000423658 | -11.2048 | *5.21E-14* |
| Unclassified_Viruses | Unclassified_Viruses | 0.000207136 | -12.2371 | *1.04E-07* |
| Uroviricota | Unclassified _Amalieviridae | 4.40563E-09 | -27.758 | *1.15E-51* |
| Phixviricota | Unclassified _Microviridae | 3.67786E-09 | -28.0185 | *3.34E-32* |
| Unclassified_Viruses | Unclassified_Viruses | 3.65981E-09 | -28.0256 | *3.34E-32* |
| Unclassified_Duplodnaviria | Unclassified_Duplodnaviria | 3.56502E-09 | -28.0634 | *1.43E-36* |
| Uroviricota | Unclassified_Caudoviricetes | 3.54907E-09 | -28.0699 | *3.66E-29* |
| Uroviricota | Unclassified_Caudoviricetes | 3.53954E-09 | -28.0738 | *8.1E-37* |
| Uroviricota | Unclassified_Caudoviricetes | 2.80063E-09 | -28.4116 | *2.16E-32* |
| Uroviricota | Unclassified _Siphoviridae | 2.6152E-09 | -28.5104 | *1.05E-24* |
| Unclassified_Duplodnaviria | Unclassified_Duplodnaviria | 2.57252E-09 | -28.5342 | *7.65E-33* |
| Uroviricota | Unclassified_Caudoviricetes | 2.54323E-09 | -28.5507 | *1.99E-29* |
| Hofneiviricota | Inoviridae_environmental samples | 2.33397E-09 | -28.6746 | *1.18E-28* |
| Unclassified_Viruses | Unclassified_Viruses | 2.15773E-09 | -28.7878 | *6.95E-35* |
| Unclassified_Viruses | Unclassified_Viruses | 1.99133E-09 | -28.9036 | *4.76E-28* |
| Uroviricota | Unclassified_Caudoviricetes | 1.83505E-09 | -29.0215 | *7.35E-50* |
| Duplornaviricota | Cystoviridae_Cystovirus | 1.37112E-09 | -29.442 | *2.24E-44* |
| Unclassified_Viruses | Unclassified_Viruses | 1.32263E-09 | -29.4939 | *1.97E-43* |
| Uroviricota | Unclassified_Caudoviricetes | 1.18786E-09 | -29.649 | *2.48E-44* |
| Uroviricota | Unclassified_Caudoviricetes | 1.18085E-09 | -29.6575 | *1.07E-39* |

| | | | | |
|---|---|---|---|---|
| Uroviricota | Unclassified_Caudoviricetes | 7.47372E-10 | -30.3175 | *1.64E-52* |
| Unclassified_Viruses | Unclassified_Viruses | 7.07462E-10 | -30.3966 | *1.19E-52* |
